# Regulation of auditory fear discrimination by the novel Kv3 voltage-gated potassium channel modulator AUT00206

**DOI:** 10.1101/2021.06.15.448180

**Authors:** Christine Stubbendorff, Ed Hale, Harriet L.L. Day, Jessica Smith, Giuseppe S. Alvaro, Charles H. Large, Carl W. Stevenson

## Abstract

Psychiatric diseases like anxiety-related disorders and schizophrenia are characterized by impaired cognition and emotional regulation linked to corticolimbic disinhibition. Restoring the balance between excitation and inhibition in corticolimbic circuits may therefore ameliorate certain features of these disorders, such as inappropriately attributing affective salience to innocuous cues. Corticolimbic activity is tightly controlled by parvalbumin-expressing GABAergic interneurons, which also regulate fear discrimination. The voltage-gated potassium channels Kv3.1 and Kv3.2 are highly expressed in these neurons, therefore Kv3.1/3.2 modulation may have potential for treating disorders associated with cognitive and emotional dysregulation. We determined the effects of the novel Kv3.1/3.2 positive modulator AUT00206 on fear discrimination. Female rats underwent limited or extended auditory fear discrimination training that we previously showed leads to discrimination or generalization, respectively, based on passive fear responding (i.e. freezing). We also assessed darting as an active fear response. We found that limited training resulted in discrimination based on freezing, which was unaffected by AUT00206. In contrast, we found that extended training resulted in generalization based on freezing and the emergence of discrimination based on darting. Importantly, AUT00206 had dissociable effects on fear discrimination and expression with extended training. While AUT00206 mitigated generalization without affecting expression based on freezing, it reduced expression without affecting discrimination based on darting. Our results indicate that Kv3.1/3.2 modulation regulates the attribution of affective significance to threat- and safety-related cues in a response-specific manner. This suggests that targeting Kv3.1 and Kv3.2 channels may provide a promising avenue for treating cognitive and emotional dysregulation in psychiatric disease.

## Introduction

Various psychiatric diseases, including anxiety-related disorders and schizophrenia, are associated with deficits in cognition and emotional regulation [1–2]. A feature of these disorders is the misattribution of affective salience to innocuous stimuli [3–4], which can be investigated by examining fear discrimination using translationally relevant preclinical models. During fear discrimination, one cue (CS+) predicts threat through its association with an aversive unconditioned stimulus (US), whereas another cue (CS−) predicts the non-occurrence of the US to signal safety. Discrimination is displayed if the CS+ elicits more fear behaviour than the CS−, whereas generalization occurs if similar fear levels are elicited by both cues [5]. Fear discrimination is a form of learned fear inhibition by the CS− and this safety signalling is impaired with the overgeneralization of fear, which is observed in schizophrenia and anxiety-related disorders [6–8]. Understanding the neurobiological basis of fear discrimination may therefore help to identify novel pharmacological targets for treating cognitive and emotional dysregulation in psychiatric disease.

Fear discrimination is regulated by functional interactions between various corticolimbic areas, including primary auditory cortex (AC), medial prefrontal cortex (mPFC), and basolateral amygdala [9–12]. Parvalbumin-expressing GABAergic interneurons (PV neurons) are crucial for orchestrating activity in this corticolimbic circuitry [13–15] and these neurons also play a key role in regulating fear discrimination [16–17]. The voltage-gated potassium channels Kv3.1 and Kv3.2 are highly expressed in PV neurons [18] and pharmacological Kv3.1/3.2 modulators that regulate fast-spiking in these neurons have recently been developed [19]. Thus Kv3.1/3.2 modulation of PV neurons may provide a novel means of rescuing corticolimbic dysfunction and, in turn, aberrant affective salience attribution in psychiatric disease.

In this study we investigated the effects of AUT00206, a novel Kv3.1/3.2 channel modulator, on auditory fear discrimination. We previously showed in female rats that limited training results in fear discrimination at retrieval, whereas extended training leads to fear generalization involving impaired safety signalling [20–21]. Thus we exploited this dual phenotype in females by determining the effects of AUT00206 on fear discrimination or generalization with limited or extended training, respectively. Our previous findings were based on freezing as a prototypical passive fear response [22] but females can also express active fear responding in the form of darting during fear conditioning and discrimination [23–24], therefore darting was also characterized here. Finally, we also determined the effects of AUT00206 on anxiety-like behaviour, locomotor activity and shock sensitivity to determine if any drug effects on fear discrimination were attributable to non-specific effects on these measures.

## Materials and methods

### Animals

Naturally cycling female Lister hooded rats (Charles River, UK) weighing 165-245 g at the beginning of the experiments were used in this study. Rats were group housed (4/cage) in individually ventilated cages on a 12 hr light/dark cycle (lights on at 8:00) with free access to food and water. All experimental procedures were conducted with ethical approval from the University of Nottingham Animal Welfare and Ethical Review Body and in accordance with the Animals (Scientific Procedures) Act 1986, UK (Home Office Project Licence 30/3230). Separate cohorts of rats were used in Experiments 1-4. All behavioral testing occurred during the animals’ light cycle.

### Drug

AUT00206 (5,5-dimethyl-3-[2-(7-methylspiro[2H-benzofuran-3,1’-cyclopropane]-4-yl)oxypyrimidin-5-yl]imidazolidine-2,4-dione) was suspended in vehicle (12.5% captisol (w/v), 0.5% hydroxypropyl methylcellulose (w/v), 0.1% Tween 80 (v/v) in distilled water) and injected (i.p.) at 30 mg/kg (in 0.2 mL/100 g). This dose was based on previous preliminary results showing that a comparable dose of AUT00206 ameliorated cognitive deficits induced by subchronic phencyclidine administration [25].

### Experiment 1: Effect of AUT00206 on auditory fear discrimination with limited training

The paradigm used for fear discrimination learning and memory testing with limited training was adapted from our previous study [20] and is illustrated in Fig 1A. Two behavioural testing chambers were used and the apparatus has been described elsewhere [26]. On Day 1, animals were habituated to two contexts (A and B), where they received two presentations each of 2 and 9 kHz tones (30 s, 80 dB, 2 min inter-trial interval (ITI)). On Day 2, animals underwent discrimination training in context A, consisting of five pairings of one tone (CS+; 30 s, 80 dB, 2 min ITI) with footshock (0.5 s, 0.4 mA, ending at tone offset) and five presentations of the other tone alone (CS−; 30 s, 80 dB, 2 min ITI). The tones used for the CS+ or CS− were counterbalanced between animals. On Day 3, animals received two presentations each of the CS+ and CS− in context B to assess memory retrieval. Tone and footshock presentations were controlled by a PC running MED-PC IV software (Med Associates, US). Animals were injected with AUT00206 or vehicle 30 min before training (Day 2) and/or retrieval (Day 3), resulting in four groups (n=9-10/group): vehicle given before training and retrieval (VEH/VEH), vehicle given before training and AUT00206 given before retrieval (VEH/AUT), AUT00206 given before training and vehicle given before retrieval (AUT/VEH), and AUT00206 given before training and retrieval (AUT/AUT). Behaviour on Days 2-3 was recorded for later data analysis.

**Fig 1.**
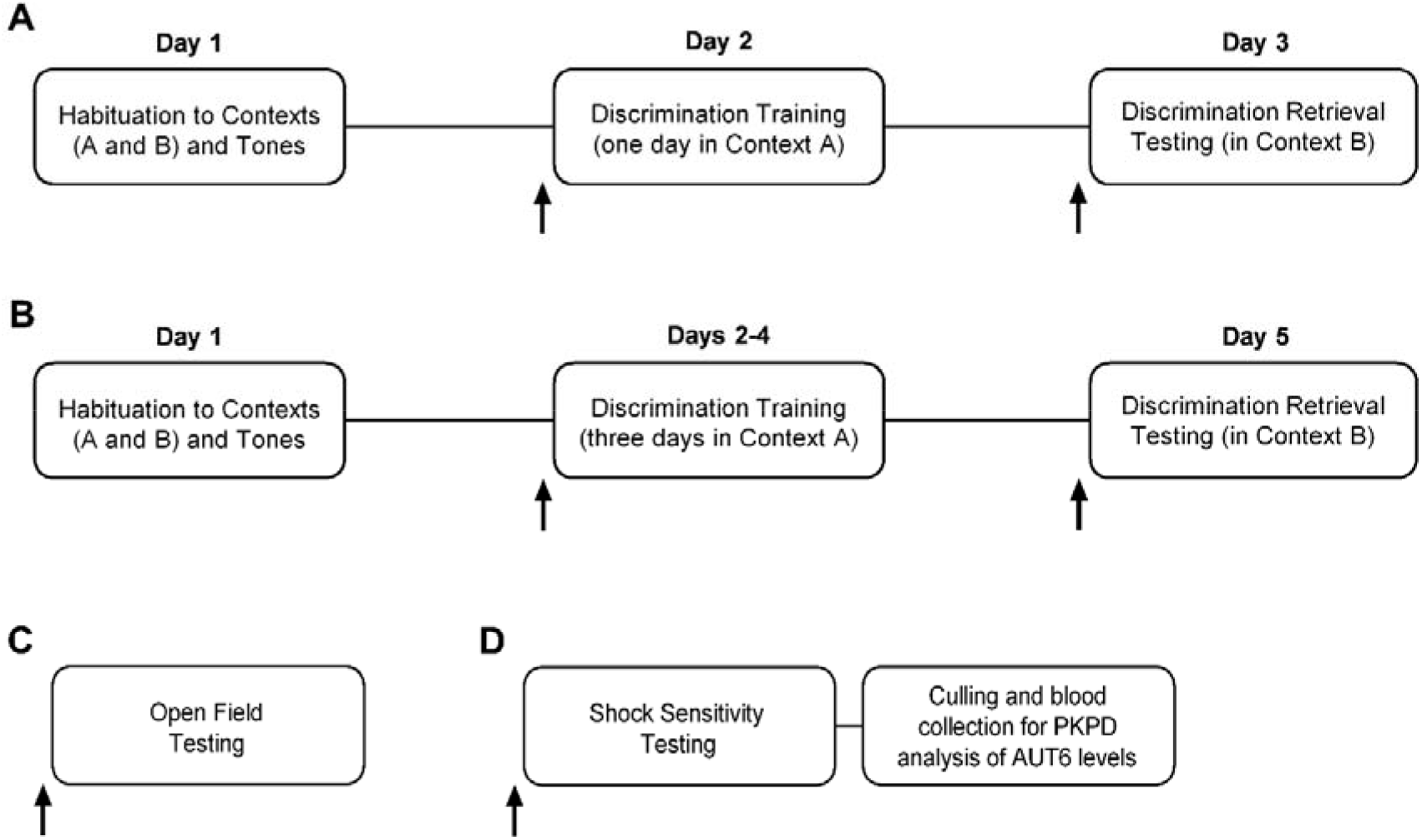
Schematic representation of the behavioural procedures used in Experiments 1-4. A) The limited discrimination training paradigm used in Experiment 1. B) The extended discrimination training paradigm used in Experiment 2. C) Animals underwent open field testing in Experiment 3. D) Animals underwent shock sensitivity testing and immediately afterwards were culled and blood was collected for the later quantification of blood AUT00206 concentration in Experiment 4.

### Experiment 2: Effect of AUT00206 on auditory fear discrimination with extended training

The paradigm used for discrimination learning and retrieval with extended training was based on our previous studies [20–21] and is depicted in Fig 1B. On Day 1, animals underwent context and tone habituation as in Experiment 1. On Days 2-4, animals underwent once-daily training sessions in context A, consisting of five CS+/US pairings and five CS− presentations as in Experiment 1 except that 0.5 mA footshocks were used. On Day 5, animals underwent retrieval testing in context B as in Experiment 1. Animals (n=9-10/group) were injected with AUT00206 or vehicle 30 min before training (Days 2-4) and/or testing (Day 5), resulting in the four groups (n=9-10/group) as in Experiment 1. Behaviour on Days 2-5 was recorded for later data analysis.

### Experiment 3: Effect of AUT00206 on behaviour during open field testing

The effect of AUT00206 on behaviour in the open field test (Fig 1C) was examined as we have described elsewhere [20]. Rats were injected with AUT00206 or vehicle (n=10/group) 30 min before being placed in the open field for 10 min. Behaviour was recorded during testing for later data analysis.

### Experiment 4: Effects of AUT00206 on shock sensitivity and blood AUT00206 levels

The effect of AUT00206 on shock sensitivity (Fig 1D) was examined as we have previously described [20]. Animals were injected with AUT00206 or vehicle (n=10/group) 30 min before receiving 10 footshocks of increasing intensity (0.05-0.5 mA, 0.5 s, 1 min ITI). Behaviour was recorded during testing for data analysis. Upon completion of shock sensitivity testing, animals treated with AUT00206 were deeply anesthetized with isoflurane before blood was collected via cardiac puncture, immediately after which they were culled. The time from AUT00206 injection to blood collection was 45-60 min. For each animal, two 70 μL blood samples were placed in two tubes both containing 130 μL of 0.1 M HEPES buffered saline and stored at −20 °C. AUT00206 levels were later determined in these samples using liquid chromatography-mass spectrometry as we have previously described [27].

### Data analysis

In Experiments 1-2, freezing and darting were quantified as passive and active fear responses, respectively. Each behaviour was scored manually by a pair of trained observers blind to drug treatment. Behaviours were scored at 3 s intervals and their cumulative durations during the 30 s cue presentations were calculated and expressed as percentages of the cue duration. Although darting has previously been expressed as the dart rate or number [23–24], we expressed darting as a percentage of cue duration to facilitate direct comparisons with freezing. Fear discrimination during training was inferred from freezing and darting during each CS+/US pairing and CS− presentation. Fear discrimination during retrieval was inferred from mean freezing and darting in response to the two CS+ and two CS− presentations. Discrimination ratios calculated from mean freezing and darting in response to the CS+ and CS− during retrieval (ratio = (CS+) / (CS+ + CS−); [28]) were also determined to account for inter-individual variability in and potential drug effects on absolute freezing and darting levels. Contextual fear during retrieval was inferred from freezing and darting in the 2 min period before cue presentations and was quantified as above. For training, behavioural data from the VEH/VEH and VEH/AUT groups were combined and data from the AUT/VEH and AUT/AUT groups were combined (n=18-20/group). During training, differences between the VEH and AUT groups in response to CS+/US pairings and CS− presentations were analyzed using three-way (Experiment 1) or four-way (Experiment 2) analysis of variance (ANOVA), with treatment as the between-subject factor and CS type and trial (and day for Experiment 2) as within-subject factors. During retrieval, group differences in response to the CS+ and CS− were analyzed using three-way ANOVA, with treatment before training and treatment before retrieval as between subject factors and CS type as the within-subject factor. Discrimination ratio data were subjected to arcsine square root transformation to stabilize the variance [29] and group differences were analyzed using two-way ANOVA, with treatment before training and treatment before retrieval as between subject factors. Group differences in contextual fear were also analyzed using two-way ANOVA, with treatment before training and treatment before retrieval as between subject factors.

In Experiment 3, behaviour during open field testing was analyzed using Ethovision software (Noldus, Netherlands). The percentage of time spent in, the number of entries into, and the latency to enter the inner zone of the open field were quantified as measures of anxiety-like behaviour. The horizontal distance moved during the open field test was also quantified as a measure of locomotor activity. Group differences on these measures were analyzed using unpaired t-tests.

In Experiment 4, the threshold current eliciting flinch and vocalization responses during shock sensitivity testing were scored manually by two observers blind to drug treatment. Group differences on these measures were analyzed using two-way ANOVA, with treatment as the between-subject factor and response as the within-subject factor. Blood AUT00206 levels 45-60 min after AUT00206 injection were also quantified, which found that one of the animals had no detectable AUT00206 in its blood. The shock sensitivity data from that animal was therefore omitted from the statistical analysis.

All data are presented as the mean+SEM. All post-hoc comparisons were conducted using the Tukey’s test where indicated. The level of significance for all comparisons was set at P<0.05.

## Results

### AUT00206 has no effect on fear discrimination with limited training

The effects of AUT00206 on freezing during limited training and retrieval are shown in Fig 2A-2D. Three-way ANOVA on freezing during the one day of training revealed significant main effects of treatment (F_(1,37)_=5.93, P=0.02) and trial (F_(4,148)_=10.44, P<0.0001) but not CS (Table S1). It also revealed a significant CS x trial interaction (F_(4,148)_=4.24, P=0.003) but no other interactions (Table S1). Post-hoc analysis showed that, compared to vehicle, AUT00206 significantly increased freezing during training across all trials and both CS types (P<0.05; Fig 2A). Two-way ANOVA on freezing before cue presentations during retrieval showed no main effects of treatment before training or treatment before retrieval, and no interaction between these factors (Table S1), indicating a lack of effect of AUT00206 on contextual fear (Fig 2B). Three-way ANOVA on freezing during cue presentations at retrieval revealed significant main effects of treatment before retrieval (F_(1,35)_=7.66, P=0.009) and CS (F_(1, 35)_=20.26, P<0.0001) but not treatment before training (Table S1). There were no interactions between any of the factors (Table S1). Post-hoc analysis showed that freezing was significantly increased during the CS+, compared to the CS−, across all groups (P<0.05), indicating successful discrimination. Freezing was significantly increased in the VEH/AUT and AUT/AUT, compared to the VEH/VEH and AUT/VEH, groups across both CS types (P<0.05), indicating that AUT00206 given before retrieval enhanced freezing in response to the cues during retrieval (Fig 2C). Two-way ANOVA on the discrimination ratio based on freezing showed no main effects of treatment before training or treatment before retrieval, and no interaction between these factors (Table S1), indicating that AUT00206 had no effect on discrimination (Fig 2D).

**Fig 2.**
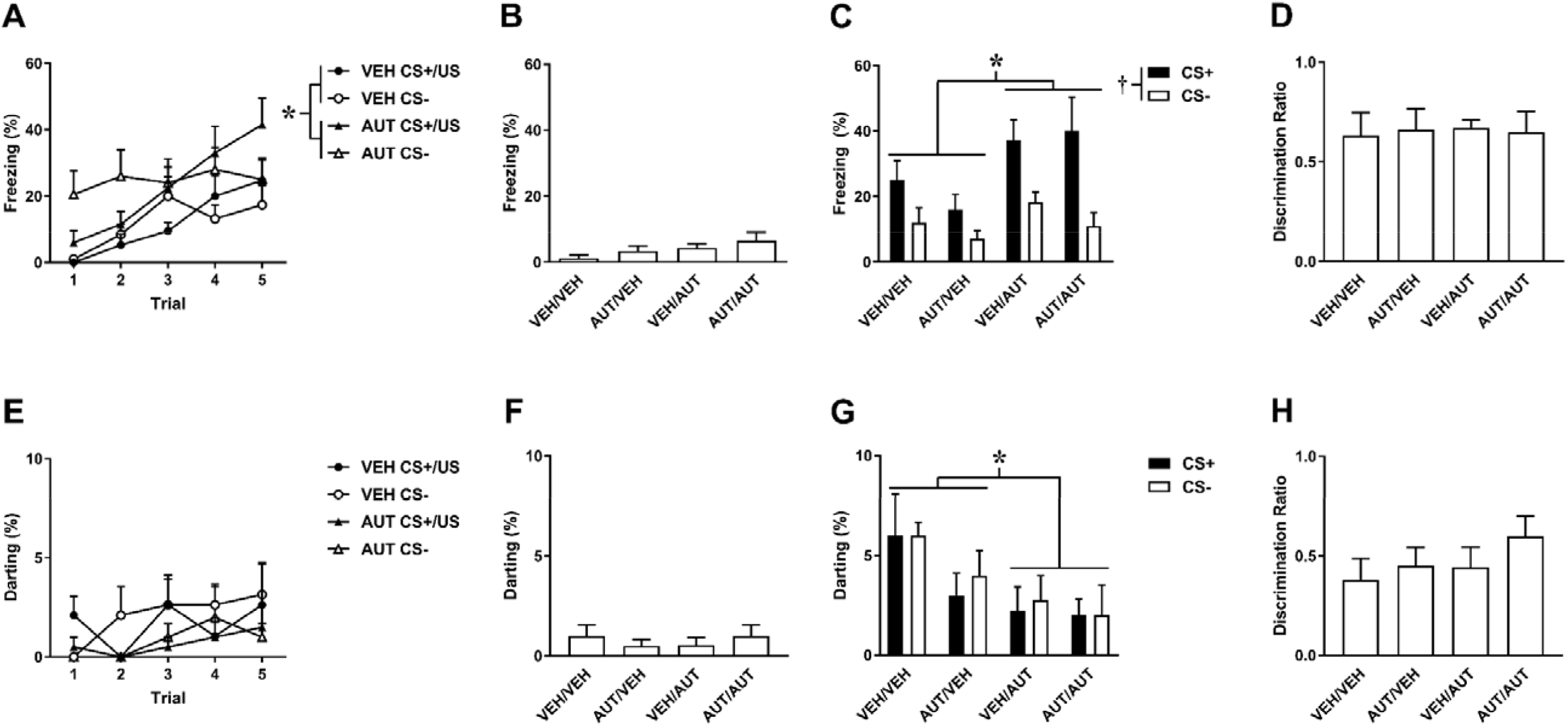
Effects of AUT00206 given before fear discrimination learning and/or memory retrieval with limited training, based on passive (A-D) and active (E-H) fear responding. A) Freezing in response to CS+/US pairings and CS− presentations during training. Freezing was increased by AUT00206, compared to vehicle (VEH), across CS+/US pairings and CS− presentations (*P<0.05). B) Freezing before cue presentations during retrieval was unaffected by AUT00206. C) Freezing in response to the CS+ and CS− at retrieval. Freezing was increased during the CS+, compared to the CS−, across all groups (^†^P<0.05). Freezing was also increased in the VEH/AUT and AUT/AUT, compared to the VEH/VEH and AUT/VEH, groups across both CS types (*P<0.05). D) The discrimination ratio, based on freezing during the CS+ and CS− at retrieval, was unaffected by AUT00206. E) Darting in response to CS+/US pairings and CS− presentations during training. AUT00206 had no effect on darting during training. F) Darting before cue presentations during retrieval was unaffected by AUT00206. G) Darting in response to the CS+ and CS− at retrieval. There were no differences in darting during the CS+, compared to the CS−, across all groups. Darting was decreased in the VEH/AUT and AUT/AUT, compared to the VEH/VEH and AUT/VEH, groups across both CS types (*P<0.05). H) The discrimination ratio, based on darting during the CS+ and CS− at retrieval, was unaffected by AUT00206 (all data presented as mean+SEM, n=9-10/group).

The effects of AUT00206 on darting during limited training and retrieval are shown in Fig 2E-2H. The levels of darting were very low throughout in comparison to freezing. Three-way ANOVA on darting during the one day of training showed no main effects of treatment, CS or trial, and no interactions between any of these factors (Table S2), indicating a lack of effect of AUT00206 on darting during training (Fig 2E). Two-way ANOVA on darting before cue presentations during retrieval showed no main effects of treatment before training or treatment before retrieval, and no interaction between these factors (Table S2), indicating a lack of effect of AUT00206 on contextual fear (Fig 2F). Three-way ANOVA on darting during cue presentations at retrieval revealed a significant main effect of treatment before retrieval (F_(1,35)_=8.03, P=0.008) but no main effects of treatment before training or CS, and no interactions between any of the factors (Table S2). Despite the low levels observed, post-hoc analysis showed that darting was significantly decreased in the VEH/AUT and AUT/AUT, compared to the VEH/VEH and AUT/VEH, groups (P<0.05), indicating that AUT00206 given before retrieval reduced darting in response to the cues during retrieval (Fig 2G). Two-way ANOVA on the discrimination ratio based on darting showed no main effects of treatment before training or treatment before retrieval, and no interaction between these factors (Table S2), indicating that AUT00206 had no effect on discrimination (Fig 2H).

### AUT00206 has dissociable response-dependent effects on fear expression and discrimination with extended training

The effects of AUT00206 on freezing during extended training and retrieval are shown in Fig 3A-3F. Four-way ANOVA on freezing during the three days of training revealed significant main effects of treatment (F_(1,35)_=5.63, P=0.023), trial (F_(4,140)_=8.00, P<0.0001), and day (F_(2,70)_=4.91, P=0.010) but not CS (Table S3). It also revealed significant CS x trial (F_(4,140)_=8.06, P<0.0001) and trial x day (F_(8,280)_=21.76, P<0.0001) interactions. There were no other interactions between the factors (Table S3). Post-hoc analysis showed that AUT00206 significantly increased freezing, compared to VEH, across all days, trials, and both CS types (P<0.05; Fig 3A-3C). Two-way ANOVA on freezing before cue presentations during retrieval revealed a significant main effect of treatment before training (F_(1,33)_=5.85, P=0.021) but no main effect of treatment before retrieval or interaction between these factors (Table S3). Post-hoc analysis showed that freezing was significantly decreased in the AUT/VEH and AUT/AUT, compared to the VEH/VEH and VEH/AUT, groups (P<0.05), indicating that AUT00206 given before training reduced contextual fear during retrieval (Fig 3D). Three-way ANOVA on freezing during cue presentations at retrieval revealed a significant main effect of CS (F_(1,33)_=10.14, P=0.003) but no main effects of treatment before training or treatment before retrieval, and no interactions between any of the factors (Table S3). Post-hoc analysis showed that freezing was significantly increased during the CS+, compared to the CS−, across all groups (P<0.05), indicating successful discrimination. Despite the lack of significant interactions, this appeared to be driven by differences between the CS+ and CS− with AUT00206 treatment as VEH/VEH controls seemed to generalize, which was confirmed by a direct comparison showing no difference in freezing between the CS+ and CS− (t_8_=0, P>0.05) in this group (Fig 3E). Two-way ANOVA on the discrimination ratio based on freezing revealed a significant main effect of treatment before training (F_(1,33)_=4.70, P=0.038) but no main effect of treatment before retrieval or interaction between these factors (Table S3). Post-hoc analysis showed that the discrimination ratio was significantly increased in the AUT/VEH and AUT/AUT, compared to the VEH/VEH and VEH/AUT, groups (P<0.05), indicating that AUT00206 given before training enhanced discrimination at retrieval based on freezing (Fig 3F).

**Fig 3.**
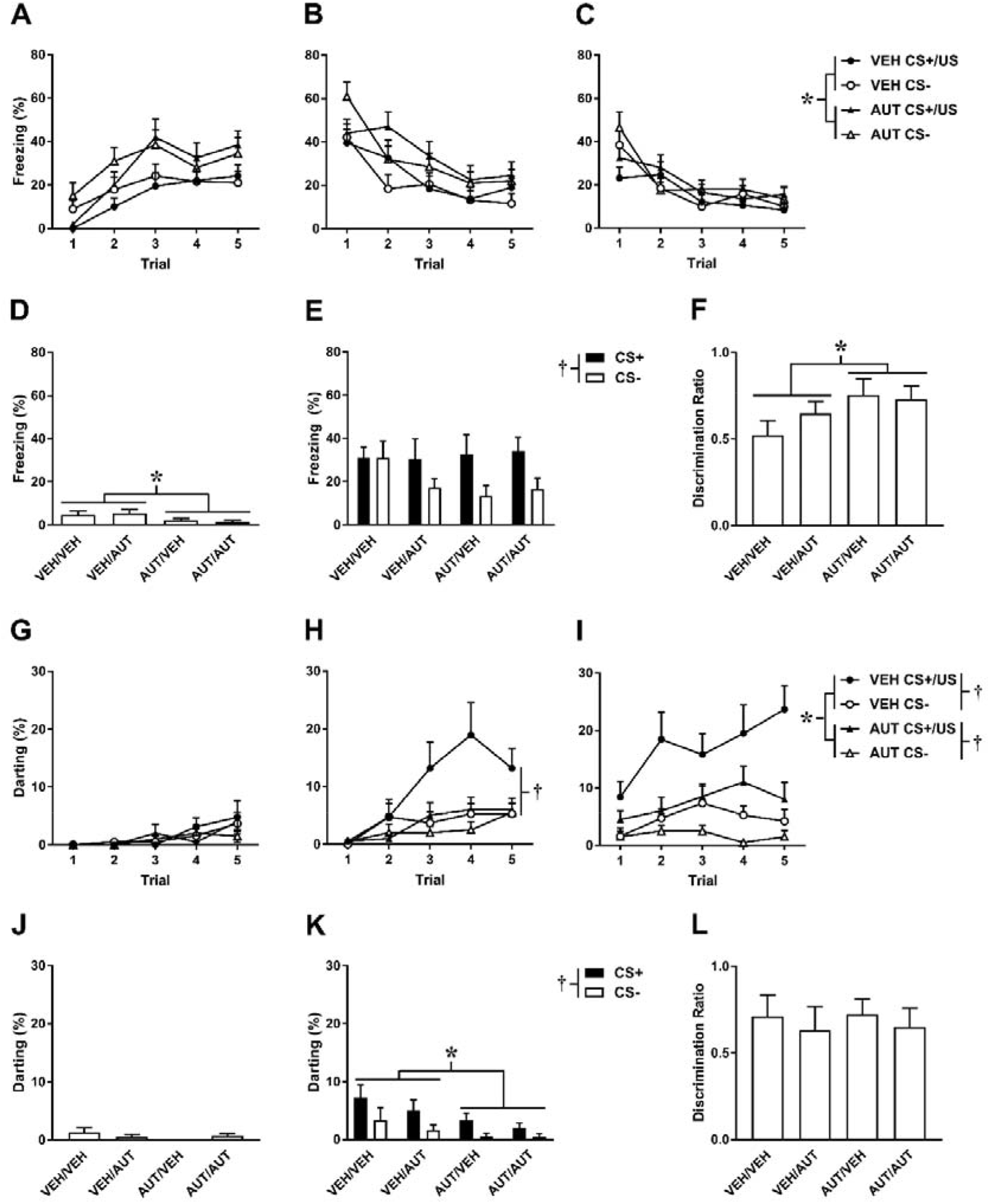
Effects of AUT00206 given before fear discrimination learning and/or memory retrieval with extended training, based on passive (A-F) and active (G-L) fear responding. A-C) Freezing in response to CS+/US pairings and CS− presentations during the first (A), second (B), and third (C) days of training. Freezing was increased by AUT00206, compared to VEH, across all days and both CS types during training (*P<0.05). D) Freezing before cue presentations during retrieval was decreased in the AUT/VEH and AUT/AUT, compared to the VEH/VEH and VEH/AUT, groups (*P<0.05). E) Freezing in response to the CS+ and CS− at retrieval. Freezing was increased during the CS+, compared to the CS−, across all groups (^†^P<0.05). F) The discrimination ratio, based on freezing during the CS+ and CS− at retrieval, was increased in the AUT/VEH and AUT/AUT, compared to the VEH/VEH and VEH/AUT, groups (*P<0.05). G-I) Darting in response to CS+/US pairings and CS− presentations during the first (G), second (H), and third (I) days of training. G) AUT00206 had no effect on darting on the first day of training. H) Darting was increased in response to the CS+/US, compared to the CS−, in the VEH (^†^P < 0.05), but not the AUT, group on the second day of training. I) On the third day of training, darting was increased in response to the CS+/US, compared to the CS−, in the VEH and AUT groups (^†^P<0.05). Darting was also decreased by AUT00206, compared to VEH, in response to the CS+/US and CS− (*P<0.05). J) Darting before cue presentations during retrieval testing was unaffected by AUT00206. K) Darting in response to the CS+ and CS− at retrieval. Darting was increased during the CS+, compared to the CS−, across all groups (^†^P<0.05). Darting was also decreased in the AUT/VEH and AUT/AUT, compared to the VEH/VEH and VEH/AUT, groups across both CS types (*P<0.05). L) The discrimination ratio, based on darting during the CS+ and CS− at retrieval, was unaffected by AUT00206 (all data presented as mean+SEM, n=9-10/group).

The effects of AUT00206 on darting during extended training and retrieval are shown in Fig 3G-3L. Four-way ANOVA on darting during the three days of training revealed significant main effects of treatment (F_(1,35)_=4.66, P=0.038), CS (F_(1,35)_=19.85, P<0.0001), trial (F_(4,140)_=12.83, P<0.0001), and day (F_(2,70)_=30.23, P<0.0001). It also revealed significant treatment x day (F_(2,70)_=7.03, P=0.002), CS x trial (F_(4,140)_=4.80, P=0.001), CS x day (F_(2,70)_=23.33, P<0.0001), trial x day (F_(8,280)_=2.28, P=0.022), treatment x CS x day (F_(2,70)_=4.26, P=0.018), and CS x trial x day (F_(8,280)_=2.34, P=0.019) interactions. There were no other interactions between the factors (Table S4). On the first day of training, darting levels were very low overall and post-hoc analysis showed that AUT00206 had no effect on darting (Fig 3G). On the second day of training, darting was significantly increased in response to the CS+/US, compared to the CS−, in the VEH (P<0.05) but not the AUT00206 group (Fig 3H). On the third day of training, all animals in the VEH group showed at least some darting with CS+/US pairings. Darting was significantly increased in response to the CS+/US, compared to the CS−, in both the VEH and AUT groups (P<0.05); darting was also significantly decreased in the AUT00206, compared to the VEH, group in response to both the CS+/US and CS− (P<0.05; Fig 3I). Darting levels before cue presentations during retrieval were very low and two-way ANOVA showed no main effects of treatment before training or treatment before retrieval, and no interaction between these factors (Table S4), indicating that AUT00206 had no effect on contextual fear before retrieval (Fig 2J). Three-way ANOVA on darting during cue presentations at retrieval revealed significant main effects of treatment before training (F_(1,33)_=8.73, P=0.006) and CS (F_(1,33)_=7.21, P=0.011) but not treatment before retrieval, and there were no interactions between any of the factors (Table S4). Post-hoc analysis showed that darting was significantly increased during the CS+, compared to the CS−, across all groups (P<0.05), indicating successful discrimination. Despite the lower levels observed in comparison to Days 2-3 of training, darting was also significantly decreased in the AUT/VEH and AUT/AUT, compared to the VEH/VEH and VEH/AUT, groups across both CS types (P<0.05), indicating that AUT00206 given before training reduced darting at retrieval (Fig 3K). Two-way ANOVA on the discrimination ratio based on darting during retrieval showed no main effects of treatment before training or treatment before retrieval, and no interaction between these factors (Table S4), indicating that AUT00206 had no effect on discrimination based on this measure (Fig 3L).

### AUT00206 reduces locomotor activity in the open field test but has no effect on shock sensitivity

The effects of AUT00206 on behaviour during open field testing are shown in Fig 4A-4D. Compared to vehicle, AUT00206 significantly decreased the time spent in (t_18_=2.79, P=0.012; Fig 4A) and number of entries into (t_18_=2.66, P=0.016; Fig 4B), but not latency to enter (t_18_=0.067, P=0.95; Fig 4C), the inner zone of the open field. AUT00206 also significantly decreased the horizontal distance moved in the open field (t_18_=2.88, P=0.01; Fig 4D), compared to vehicle, indicating that its effects were likely due to reduced locomotion rather than enhanced anxiety-like behaviour.

**Fig 4.**
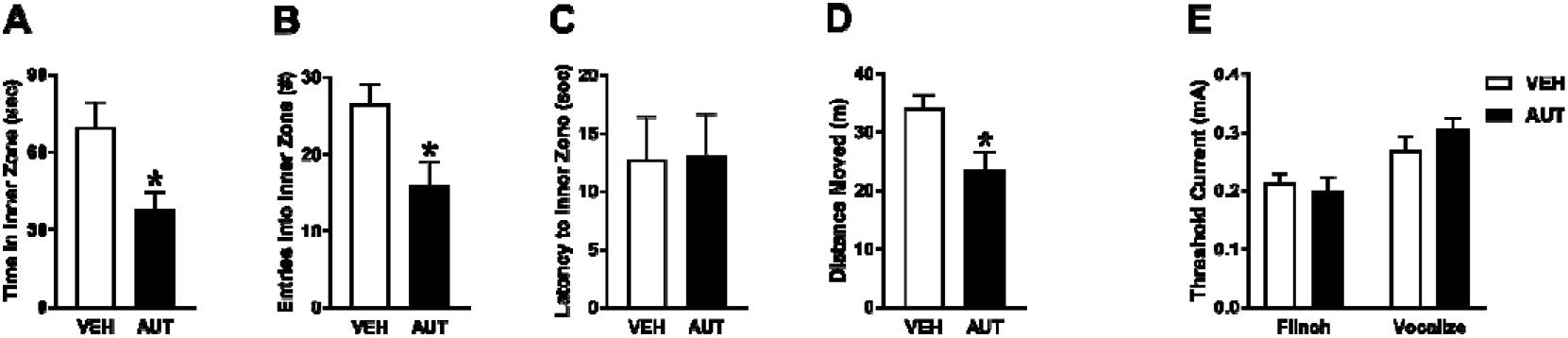
Effects of AUT00206 on behaviour during open field testing and on shock sensitivity. A-B) AUT00206 decreased the time spent in (A) and number of entries into (B) the inner zone of the open field (*P<0.05). C) AUT00206 had no effect on the latency to enter the inner zone of the open field. D) AUT00206 decreased the horizontal distance moved in the open field (*P<0.05). E) AUT00206 had no effect on the threshold current eliciting flinch responses or audible vocalizations (all data presented as mean+SEM, n=9-10/group).

The effects of AUT00206 on shock sensitivity are shown in Fig 4E. Two-way ANOVA revealed a significant main effect of response (F_(1,17)_=24.57, P=0.0001) but no main effect of treatment (F_(1,17)_=0.20, P=0.66) or treatment x response interaction (F_(1,17)_=2.44, P=0.14), indicating that AUT00206 had no effect on the threshold current eliciting flinch or vocalization responses. Mean AUT00206 concentration in the blood 45-60 min after AUT00206 treatment was 7.85±1.28 μg/mL (range=1.1-13.4 μg/mL).

## Discussion

In this study we found fear discrimination at retrieval after limited training based on freezing, while darting expression was low and no discrimination was observed with respect to this measure. AUT00206 increased freezing and decreased darting acutely but had no effect on freezing-based discrimination with limited training. In contrast, with extended training we found generalization in relation to freezing, whereas discrimination emerged based on darting. AUT00206 increased freezing and decreased darting acutely during extended training. Importantly, AUT00206 given before extended training had dissociable effects on fear expression and discrimination during later retrieval in a response-specific manner. AUT00206 enhanced discrimination without affecting fear responding to the CS+ in relation to freezing, whereas it reduced cue-induced fear responding without affecting discrimination based on darting. AUT00206 also reduced locomotor activity in the open field test but had no effect on shock sensitivity, indicating that increased freezing and decreased darting with acute AUT00206 treatment may have involved non-specific drug effects on locomotion. However, this cannot explain the lasting effects of AUT00206 given before extended training on behaviour during later retrieval. Our results instead suggest that AUT00206 mitigated fear generalization based on passive responding while also reducing fear expression based on active responding.

We found previously that whether females discriminate or generalize depends on the extent of discrimination training received [20–21]. Our results confirm these previous findings showing discrimination with limited training and generalization with extended training, based on freezing at retrieval. Here we also examined darting as this active fear response has been reported in females during fear conditioning and discrimination in some [23–24], but not all [30–32], recent studies. With limited training we found low levels of darting and no evidence of discrimination in relation to this measure. However, darting levels increased and discrimination based on darting developed over the course of extended training, in agreement with a recent study [24]. Differences in darting during limited vs extended training may reflect a switch from passive to active fear responding with greater threat imminence, such that higher darting might be expected with more CS+/US pairings. Furthermore, a shift back towards passive fear responding in the absence of the US might explain why darting returned to lower levels during later retrieval [33].

Recent optogenetic studies provide compelling evidence that cortical PV neurons are crucial for regulating fear discrimination and safety signalling as indexed by freezing. While activation of PV neurons in AC had no effect on auditory fear discrimination, their inactivation caused generalization [16]. In mPFC, PV neurons bidirectionally regulate auditory fear discrimination. During discrimination training, PV neuronal firing was associated with safety signalling by the CS−. Moreover, PV neuronal activation enhanced discrimination, whereas inactivation of PV neurons resulted in generalization involving reduced safety signalling by the CS− [17]. Kv3.1 and Kv3.2 channels are highly expressed on cortical PV neurons and activation of these channels is important for maintaining their fast-firing phenotype [18]. In the present study AUT00206 had no effect on discrimination with limited training but it did prevent generalization from developing with extended training. Interestingly, a recent study showed that Kv3.1/3.2 modulation had no effect on cortical PV neuronal firing when given alone but it did reverse the pharmacological perturbation of activity in these neurons [19]. This raises the intriguing possibility that extended discrimination training resulted in fear generalization by altering PV neuronal function, which was rescued by AUT00206, although this remains to be determined in future studies.

We showed that AUT00206 treatment during extended training also dampened cue-induced darting at later retrieval independently of any effect on discrimination based on this measure. Although the neurochemical mechanisms underpinning darting have begun to be identified [34], the neural circuit basis of darting still needs to be elucidated. The activation of parvalbumin-expressing projection neurons in different corticothalamic, corticostriatal, and tectofugal circuits mediates various active fear behaviors [35–39]. However, we found that AUT00206 reduced darting on the last day of extended training and had no acute effect on this active fear response at retrieval. Instead we found that darting during retrieval was reduced by prior AUT00206 treatment during extended training. Taken together, these results suggest that other circuit mechanisms might underpin the regulation of darting by PV neurons.

## Conclusion

This study is the first to show that pharmacologically modulating Kv3.1 and Kv3.2 channels regulates fear discrimination and expression in a response-specific manner. This suggests that Kv3.1/3.2 channel modulation may provide a novel target for treating cognitive and emotional dysregulation in psychiatric disease, such as the misattribution of emotional salience to harmless environmental cues. Kv3.1 and Kv3.2 channels are highly expressed by PV neurons, which are crucial for modulating corticolimbic circuit function. Importantly, diseases such as anxiety-related disorders and schizophrenia are associated with corticolimbic circuit dysfunction, which might be ameliorated by Kv3.1/3.2 modulators given alone or as adjuncts to other first-line treatments. This study also highlights the utility of characterizing both passive and active fear responding to gain a fuller appreciation of the behavioural basis of fear discrimination, although further research is required to characterize the neurobiological mechanisms underlying the darting response.

## Funding and Disclosure

This study was funded by Autifony Therapeutics. CS and EH were supported by the Biotechnology and Biological Sciences Research Council (BBSRC) [grant number BB/P001149/1]. HLLD was supported by a BBSRC Doctoral Training Partnership [grant number BB/J014508/1]. JS was supported by a Wellcome Trust Biomedical Vacation Scholarship [grant number 208669/Z/17/Z]. GSA and CHL are shareholders and full-time employees of Autifony Therapeutics. The other authors declare no competing interests.

## Acknowledgments

The authors thank Federica Bianchi at Aptuit for conducting the liquid chromatography-mass spectrometry analysis of blood AUT00206 levels and Helen Cassaday for providing insightful comments on the manuscript.

## Author Contributions

CS and EH conducted the open field and shock sensitivity experiments and collected the post-mortem blood samples. CS and HLLD quantified freezing behavior. EH and JS quantified darting behavior. GSA and CHL provided essential reagents. CHL and CWS designed the study. CWS conducted the fear discrimination experiments, analyzed the data, and wrote the manuscript.

---

**Table S1.**
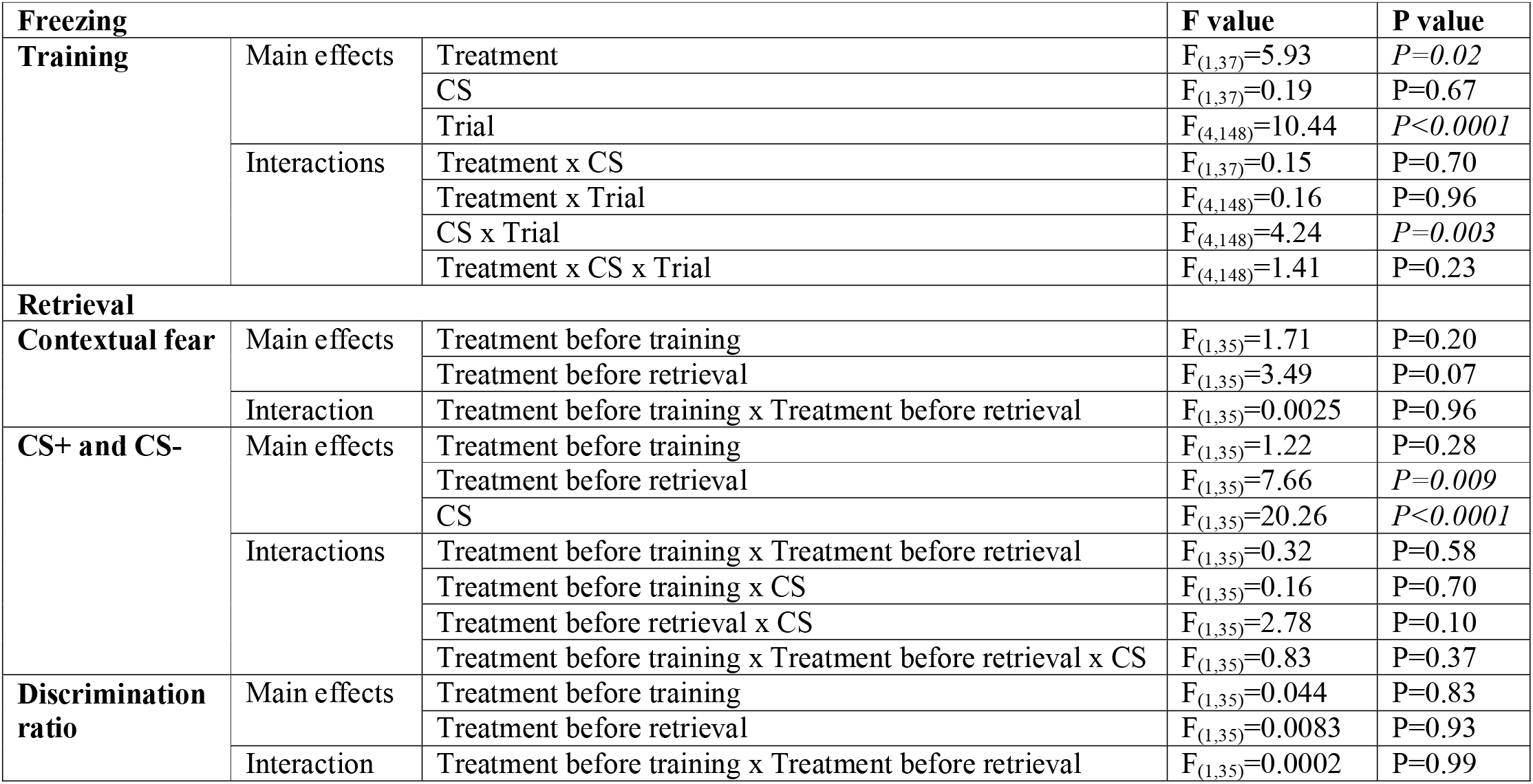
Summary of statistical analysis of freezing data from Experiment 1 (significant P values in italics)

**Table S2.**
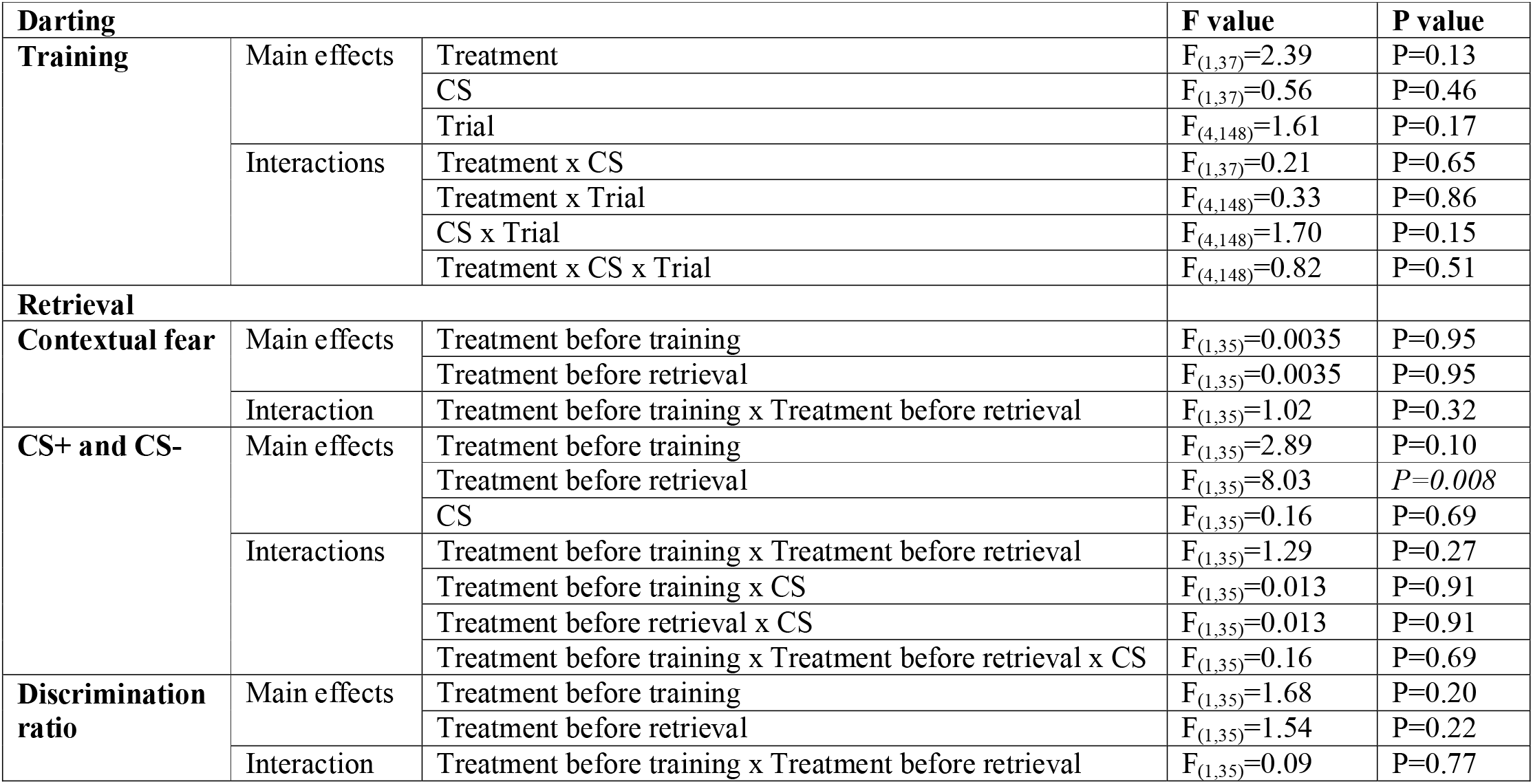
Summary of statistical analysis of darting data from Experiment 1 (significant P values in italics)

| Darting |  |  | F value | P value |
| --- | --- | --- | --- | --- |
| Training | Main effects | Treatment | $F_{(1,37)}=2.39$ | P=0.13 |
| | | CS | $F_{(1,37)}=0.56$ | P=0.46 |
| | | Trial | $F_{(4,148)}=1.61$ | P=0.17 |
| | Interactions | Treatment x CS | $F_{(1,37)}=0.21$ | P=0.65 |
| | | Treatment x Trial | $F_{(4,148)}=0.33$ | P=0.86 |
| | | CS x Trial | $F_{(4,148)}=1.70$ | P=0.15 |
| | | Treatment x CS x Trial | $F_{(4,148)}=0.82$ | P=0.51 |
| Retrieval |  |  |  |  |
| Contextual fear | Main effects | Treatment before training | $F_{(1,35)}=0.0035$ | P=0.95 |
| | | Treatment before retrieval | $F_{(1,35)}=0.0035$ | P=0.95 |
| | Interaction | Treatment before training x Treatment before retrieval | $F_{(1,35)}=1.02$ | P=0.32 |
| CS+ and CS- | Main effects | Treatment before training | $F_{(1,35)}=2.89$ | P=0.10 |
| | | Treatment before retrieval | $F_{(1,35)}=8.03$ | $P=0.008$ |
| | | CS | $F_{(1,35)}=0.16$ | P=0.69 |
| | Interactions | Treatment before training x Treatment before retrieval | $F_{(1,35)}=1.29$ | P=0.27 |
| | | Treatment before training x CS | $F_{(1,35)}=0.013$ | P=0.91 |
| | | Treatment before retrieval x CS | $F_{(1,35)}=0.013$ | P=0.91 |
| | | Treatment before training x Treatment before retrieval x CS | $F_{(1,35)}=0.16$ | P=0.69 |
| Discrimination ratio | Main effects | Treatment before training | $F_{(1,35)}=1.68$ | P=0.20 |
| | | Treatment before retrieval | $F_{(1,35)}=1.54$ | P=0.22 |
| | Interaction | Treatment before training x Treatment before retrieval | $F_{(1,35)}=0.09$ | P=0.77 |

**Table S3.** Summary of statistical analysis of freezing data from Experiment 2 (significant P values in italics)

| Freezing |  |  | F value | P value |
| --- | --- | --- | --- | --- |
| Training | Main effects | Treatment | $F_{(1,35)}=5.63$ | $P=0.023$ |
| | | CS | $F_{(1,35)}=0.56$ | $P=0.46$ |
| | | Trial | $F_{(4,140)}=8.00$ | $P<0.0001$ |
| | | Day | $F_{(2,70)}=4.91$ | $P=0.01$ |
| | Interactions | Treatment x CS | $F_{(1,35)}=0.031$ | $P=0.86$ |
| | | Treatment x Trial | $F_{(4,140)}=0.32$ | $P=0.87$ |
| | | Treatment x Day | $F_{(2,70)}=0.66$ | $P=0.52$ |
| | | CS x Trial | $F_{(4,140)}=8.06$ | $P<0.0001$ |
| | | CS x Day | $F_{(2,70)}=0.98$ | $P=0.38$ |
| | | Trial x Day | $F_{(8,280)}=21.76$ | $P<0.0001$ |
| | | Treatment x CS x Trial | $F_{(4,140)}=0.56$ | $P=0.70$ |
| | | Treatment x CS x Day | $F_{(2,70)}=0.27$ | $P=.076$ |
| | | Treatment x Trial x Day | $F_{(8,280)}=0.92$ | $P=0.50$ |
| | | CS x Trial x Day | $F_{(8,280)}=1.85$ | $P=0.067$ |
| | | Treatment x CS x Trial x Day | $F_{(8,280)}=0.35$ | $P=0.95$ |
| Retrieval |  |  |  |  |
| Contextual fear | Main effects | Treatment before training | $F_{(1,33)}=5.85$ | $P=0.021$ |
| | | Treatment before retrieval | $F_{(1,33)}=0.0091$ | $P=0.92$ |
| | Interaction | Treatment before training x treatment before retrieval | $F_{(1,33)}=0.46$ | $P=0.50$ |
| CS+ and CS- | Main effects | Treatment before training | $F_{(1,33)}=0.40$ | $P=0.53$ |
| | | Treatment before retrieval | $F_{(1,33)}=0.23$ | $P=0.64$ |
| | | CS | $F_{(1,33)}=10.14$ | $P=0.003$ |
| | Interactions | Treatment before training x treatment before retrieval | $F_{(1,33)}=0.79$ | $P=0.38$ |
| | | Treatment before training x CS | $F_{(1,33)}=2.24$ | $P=0.14$ |
| | | Treatment before retrieval x CS | $F_{(1,33)}=0.52$ | $P=0.48$ |
| | | Treatment before training x treatment before retrieval x CS | $F_{(1,33)}=0.94$ | $P=0.34$ |
| Discrimination ratio | Main effects | Treatment before training | $F_{(1,33)}=4.70$ | $P=0.038$ |
| | | Treatment before retrieval | $F_{(1,33)}=0.16$ | $P=0.69$ |
| | Interaction | Treatment before training x treatment before retrieval | $F_{(1,33)}=0.92$ | $P=0.34$ |

**Table S4.**
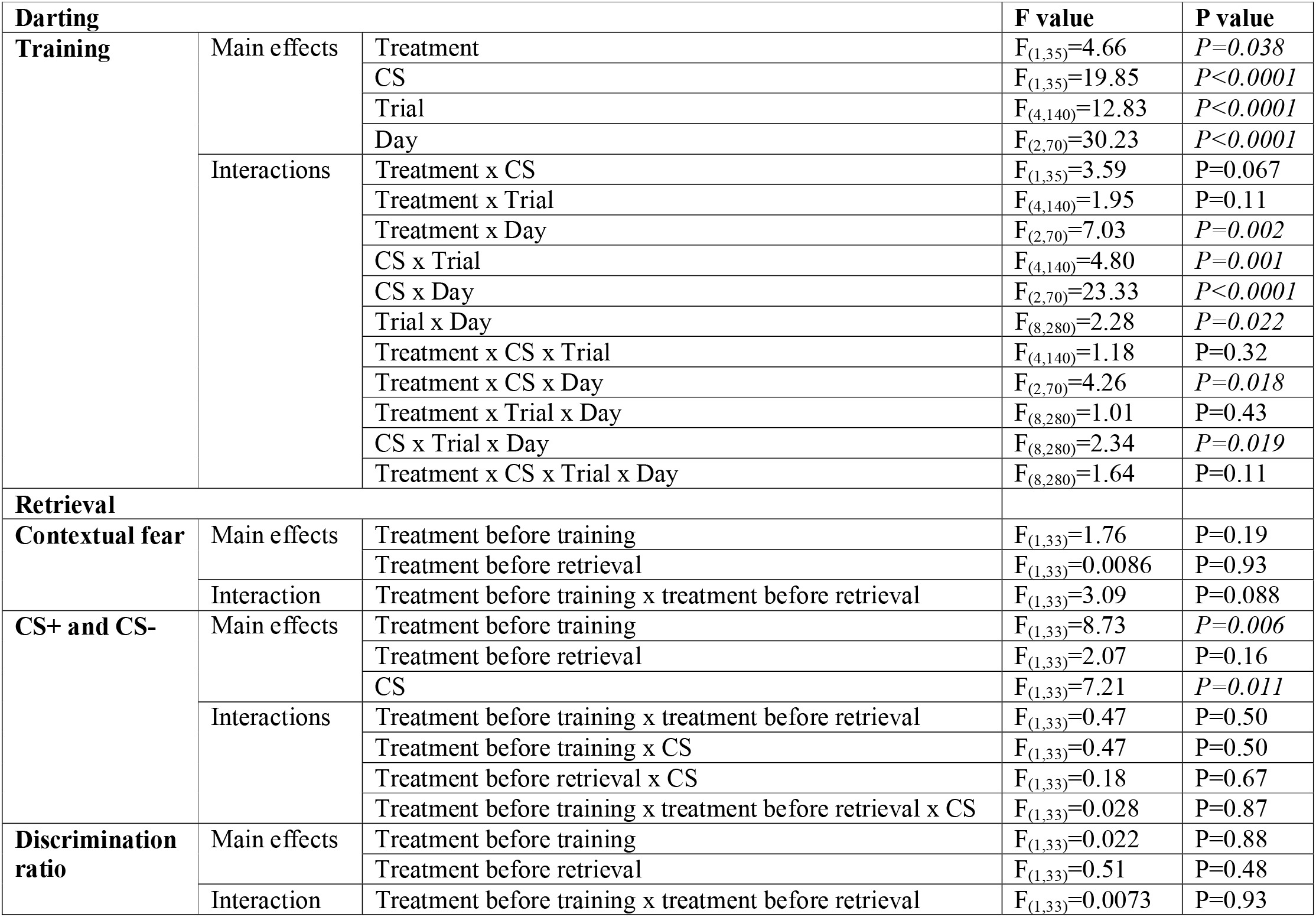
Summary of statistical analysis of darting data from Experiment 2 (significant P values in italics)

| Darting |  |  | F value | P value |
| --- | --- | --- | --- | --- |
| Training | Main effects | Treatment | $F_{(1,35)}=4.66$ | $P=0.038$ |
| | | CS | $F_{(1,35)}=19.85$ | $P<0.0001$ |
| | | Trial | $F_{(4,140)}=12.83$ | $P<0.0001$ |
| | | Day | $F_{(2,70)}=30.23$ | $P<0.0001$ |
| | Interactions | Treatment x CS | $F_{(1,35)}=3.59$ | $P=0.067$ |
| | | Treatment x Trial | $F_{(4,140)}=1.95$ | $P=0.11$ |
| | | Treatment x Day | $F_{(2,70)}=7.03$ | $P=0.002$ |
| | | CS x Trial | $F_{(4,140)}=4.80$ | $P=0.001$ |
| | | CS x Day | $F_{(2,70)}=23.33$ | $P<0.0001$ |
| | | Trial x Day | $F_{(8,280)}=2.28$ | $P=0.022$ |
| | | Treatment x CS x Trial | $F_{(4,140)}=1.18$ | $P=0.32$ |
| | | Treatment x CS x Day | $F_{(2,70)}=4.26$ | $P=0.018$ |
| | | Treatment x Trial x Day | $F_{(8,280)}=1.01$ | $P=0.43$ |
| | | CS x Trial x Day | $F_{(8,280)}=2.34$ | $P=0.019$ |
| | | Treatment x CS x Trial x Day | $F_{(8,280)}=1.64$ | $P=0.11$ |
| Retrieval |  |  |  |  |
| Contextual fear | Main effects | Treatment before training | $F_{(1,33)}=1.76$ | $P=0.19$ |
| | | Treatment before retrieval | $F_{(1,33)}=0.0086$ | $P=0.93$ |
| | Interaction | Treatment before training x treatment before retrieval | $F_{(1,33)}=3.09$ | $P=0.088$ |
| CS+ and CS- | Main effects | Treatment before training | $F_{(1,33)}=8.73$ | $P=0.006$ |
| | | Treatment before retrieval | $F_{(1,33)}=2.07$ | $P=0.16$ |
| | | CS | $F_{(1,33)}=7.21$ | $P=0.011$ |
| | Interactions | Treatment before training x treatment before retrieval | $F_{(1,33)}=0.47$ | $P=0.50$ |
| | | Treatment before training x CS | $F_{(1,33)}=0.47$ | $P=0.50$ |
| | | Treatment before retrieval x CS | $F_{(1,33)}=0.18$ | $P=0.67$ |
| | | Treatment before training x treatment before retrieval x CS | $F_{(1,33)}=0.028$ | $P=0.87$ |
| Discrimination ratio | Main effects | Treatment before training | $F_{(1,33)}=0.022$ | $P=0.88$ |
| | | Treatment before retrieval | $F_{(1,33)}=0.51$ | $P=0.48$ |
| | Interaction | Treatment before training x treatment before retrieval | $F_{(1,33)}=0.0073$ | $P=0.93$ |

## Notes

### Competing Interest Statement

This study was funded by Autifony Therapeutics. GSA and CHL are shareholders and full-time employees of Autifony Therapeutics.

